# Smelling sensations: olfactory crossmodal correspondences

**DOI:** 10.1101/2020.04.15.042630

**Authors:** Ryan J. Ward, Sophie M. Wuerger, Alan Marshall

## Abstract

Crossmodal correspondences are the associations between apparently distinct stimuli in different sensory modalities. These associations, albeit surprising, are generally shared in most of the population. Olfaction is ingrained in the fabric of our daily life and constitutes an integral part of our perceptual reality, with olfaction being more commonly used in the entertainment and analytical domains, it is crucial to uncover the robust correspondences underlying common aromatic compounds. Towards this end, we investigated an aggregate of crossmodal correspondences between ten olfactory stimuli and other modalities (angularity of shapes, smoothness of texture, pleasantness, pitch, colours, musical genres and emotional dimensions) using a large sample of 68 observers. We uncover the correspondences between these modalities and extent of these associations with respect to the explicit knowledge of the respective aromatic compound. The results revealed the robustness of prior studies, as well as, contributions towards olfactory integration between an aggregate of other dimensions. The knowledge of an odour’s identity coupled with the multisensory perception of the odours indicates that these associations, for the most part, are relatively robust and do not rely on explicit knowledge of the odour. Through principal component analysis of the perceptual ratings, new cross-model mediations have been uncovered between odours and their intercorrelated sensory dimensions. Our results demonstrate a collective of associations between olfaction and other dimensions, potential cross modal mediations via exploratory factor analysis and the robustness of these correspondence with respect to the explicit knowledge of an odour. We anticipate the findings reported in this paper could be used as a psychophysical framework aiding in a collective of applications ranging from olfaction enhanced multimedia to marketing.

## Introduction

Olfaction is ingrained into the fabric of our lives, altering the very perception of our favourite commodities and plays a crucial role in the multi-sensory perception of our surrounding environment. Exploring the crossmodal correspondences provided by the chemical senses is pivotal towards solving the crossmodal binding problem, and in turn, their implication on interactive and immersive experiences. Crossmodal correspondences can be depicted as the consistent correspondence between stimulus features in different sensory modalities [1]. These correspondences can be matched if they both have the same effect on the observers’ mood, emotional state, alertness and/or arousal [1–3]. Aromas are consistently perceived together with other stimuli principally visual, the absence of these additional stimuli results in inferior identification [4,5]. Albeit, semantic congruency can enhance the perceived pleasantness [6,7], discrimination [8] and correct identification of odours [9]. Crossmodal correspondences have been shown to induce a bias (i.e., providing a red glass of white wine can bias the judgment of expert wine tasters [10]). In terms of evolution, olfaction is one of the oldest senses and plays a major role in social behaviour, communication and emotional evaluation; mood and emotional processes share a common neural substrate with the olfactory pathway, namely the limbic system [11]. It is therefore likely that olfactory information plays a major role in modulating the quality of our immersive multisensorial experiences.

Crossmodal interactions between smell and both vision and hearing have been of distinct interest, as it has been shown to alter olfactory perception considerably. For example, olfaction-audition [12–14], olfaction-colour [15–18], olfaction-visual motion [19], and olfaction-angularity of shapes [18,20]. The mechanisms underlying such correspondences has diverse characterisation within the literature. The most frequently deducted mechanisms are; hedonics [14,20–23], semantics [1,18,22,24– 27] and natural co-occurrence [1,27,28]. Identifying robust crossmodal olfactory associations has a vast array of practical implications spanning from advertising to human-computer interaction. A contributory factor to idiosyncratic, as opposed to robust associations, is the relatively high inter-observer variability of the olfactory sense [29]. There is an increasing demand to provide the chemical senses to human-machine interfaces, as modern technology is typically limited to vision, auditory and basic haptic feedback. Understanding how the chemical senses interact with the other senses and their implication on interactive and immersive experiences is the next step towards total sensory immersion and augmenting artificial synaesthesia to expand and create new senses for humans.

Synaesthesia is a neurological condition in which input into one modality elicits a simultaneous perception in a second modality [30]. Controversially, synaesthesia could be considered a more severe manifestation of crossmodal associations [26] and antagonistically [31]. It is hypothesized that everyone is born with synaesthesia [32] but is later disinhibited [33] and/or pruned. Synaesthesia can still provide valuable information on the initial hypothesis between the strong intercorrelated sensory dimensions and their potential in cross modal correspondences despite the automatic evocation.

We hypothesize that there will be robust associations that are mediated predominantly by hedonic ratings underlying common aromatic compounds. These associations could, at least in part, be affected by knowledge of the odour’s identity. We provide evidence to support both hypotheses, we explored a series of potential crossmodal correspondences. Here we present associations between olfactory stimuli and the angularity of shapes, colour, smoothness of textures, pitch, musical genres, and several emotional dimensions. We further investigate whether explicit knowledge of the odours modulates these associations by using an odour identification task. Exploratory factor analysis was then conducted on the perceptual ratings, revealing potential crossmodal mediations and the perceptual similarity between the ten aromatic compounds used in these experiments.

## Materials & methods

### Participants

68 individuals (23 males and 45 females with a mean age of 26.75 (standard deviation: 12.75)) took part in the experiments. No participants reported any impairment that could affect their sense of smell (i.e., cold or flu). Participants were briefed about potential allergens and breaks (a minimum of a 10-minute break halfway through, or if the participant felt like they have got a reduced sense of smell). The experiment was given ethical approval by the University of Liverpool, lasted approximately 50 minutes and conducted in accordance to the standards set in the Declaration of Helsinki for Medical Research Involving Human Subjects. Participants gave written informed consent before taking part in the experiment.

### Apparatus

All results were obtained through a graphical user interface programmed in MATLAB R2018b. Participants were placed in a lightproof anechoic chamber equipped with an overhead luminaire (GLE-M5/32; GTI Graphic Technology Inc., Newburgh, NY) during the experiment. The lighting in the room was kept consistent by using the daylight simulator of the overhead luminaire. The speakers were JBL Desktop speakers; the colour stimuli were shown on a calibrated EIZO ColorEdge CG243W monitor.

### Tasks and Stimuli

Participants were instructed to associate a given odour with a value along each of the following dimensions: visual shapes, textures, pleasantness (using a l scale), pitch, music genre and emotions. The aromas were presented in a random order (determined by a random number generator in MATLAB) and all associations were assessed in the same order for a given aroma. At the end of the experiment, participants were asked to identify the odour (identification task). For three of the experimental tests (visual shapes, texture, pleasantness) and for the emotion task a neural option was available, but participants were strongly discouraged to use this option.

### Odour Stimuli

Ten odourants were used; five from Mystic Moments™; caramel, cherry, coffee, freshly cut grass, and pine; five from Miaroma™; black pepper, lavender, lemon, orange and peppermint. These aromas were selected as they can be frequently found during everyday life and gives diversity in the chemical makeup of the aromas. For consistency and to avoid any other associations that would affect the results of this experiment 4 mL of the respective essential oil was placed in a clear test tube, wrapped in white tape and numbered 1 through 10 in an initial random permutation. The aromas were stored at ≈ 2.5°C to minimize oxidation, all odours were removed and placed back into the fridge at the same time to ensure approximately uniform evaporation. The odours were replaced every two weeks.

### Shape Stimuli

A nine-point scale was constructed with a rounded shape “bouba” and an angular shape “kiki” on the left and right side of the scale respectively. Similar to an earlier experiment performed by [20]. The midpoint of the nine-point scale (5) was neutral (no opinion).

### Texture Stimuli

A nine-point scale was constructed with smooth and rough anchored on the left and right side respectively. Participants were supplied with representative textures to aid them in their decision, with silk being a representative for smooth and sandpaper being a representative for rough. The midpoint of the nine-point scale (5) was neutral (no opinion). Participants felt the texture at least once during the questions’ first appearance.

### Pleasantness

A nine-point scale from very unpleasant to very pleasant was used, with 5 being the neutral option.

### Pitch Stimuli

The full range of audible frequencies (20Hz to 20kHz) was implemented using a slider where movement from left to right corresponded to an increase in frequency. Every time the slider was adjusted the respective frequency was played, producing a sinusoidal tone lasting 1 second in length. Due to the large volume of potential selections, participants were played a sample from each end of the scale, followed by a sample at 10kHz, if the current pitch didn’t match the odour a lower or higher pitch was selected (approximately half way between the last two frequencies played) as indicated by the participant.

### Music Stimuli

Seven different music genres; classical, country, heavy metal, jazz, rap, classic rock, and soul. Five were selected from [34] with an additional two added due to their wide popularity. Each sample was 15 seconds in duration and played at the same volume across participants. Participants had to listen to each sample at least once during the questions first occurrence, the order was subject to the participants preference.

### Colour Stimuli

The CIE L*a*b* colour space was used because of its perceptual uniformity, participants could slide through 101 linear interpolated slices from the L* channel of the colour space increasing or decreasing the lightness. Only colours that fit in sRGB colour gamut were shown. This removed the limitations of earlier studies that let participants choose from a small selection of colours.

### Emotions

A subset of emotions from the Universal Emotion and Odour Scale [9] was included these where; angry, aroused, bored, calm, disgust, excited, happy, sad and scared, additionally an option for neutral (no opinion) was added to negate a tentative assignment.

### Identification Task

A list of different aromas was compiled consisting of the ten odours used in this experiment along with an additional fifteen and presented in alphabetical order. This was incorporated to increase the number of choices the participant had available, so they were less likely to make inferred decisions when trying to identify the current odour. The identification task was at the end of the experiment.

## Results

### Angularity, smoothness, pleasantness and pitch

Fig 1 shows the mean ratings (transformed to z-scores) for angularity, smoothness, pleasantness and pitch for each of the 10 odours. Friedman tests were conducted on the z-scores of the angularity, smoothness, pleasantness and pitch ratings to test if the odours influenced the ratings. This revealed that the odours significantly affected all ratings; angularity (*x*^2^(9) = 122.15, *P* < 0.05), smoothness (*x*^2^(9) = 52.32, *P* < 0.05), pleasantness (*x*^2^(9) = 81.85, *P* < 0.05) and pitch (*x*^2^(9) = 101.75, *P* < 0.05). A Bonferroni multiple comparison test was conducted to identify which odours where significantly different from each other. A post-hoc one sample t-test with a Bonferroni correct alpha was conducted to determine which of the odours was significantly different from 0 (the original scale’s grand mean). The significantly ‘rounded’ odours are caramel (*P* < 0.005, *t* = -9.88) and coffee (*P* < 0.005, *t* = -3.87). The significantly ‘angular’ odours are peppermint (*P* < 0.005, *t* = 8.62) and lemon (*P* < 0.005, *t* = 3.43), as shown in Fig 1A. The significantly ‘rough’ odour is black pepper (*P* < 0.005, *t* = -3.22) the significantly ‘smooth’ odour is caramel (*P* < 0.005, *t* = 4.64), as shown in Fig 1B. The significantly ‘pleasant’ odours are lemon (*P* < 0.005, *t* = 4.27) and orange (*P* < 0.005, *t* = 6.87). The significantly ‘unpleasant’ odour is black pepper (*P* < 0.005, *t* = -5.84), as shown in Fig 1C. The significantly ‘higher pitch’ odour is peppermint (*P* < 0.005, *t* = 3.47). The significantly ‘lower pitch’ odours are coffee (*P* < 0.005, *t* = -5.64) and caramel (*P* < 0.005, *t* = -4.60), as shown in Fig 1D.

**Fig 1.**
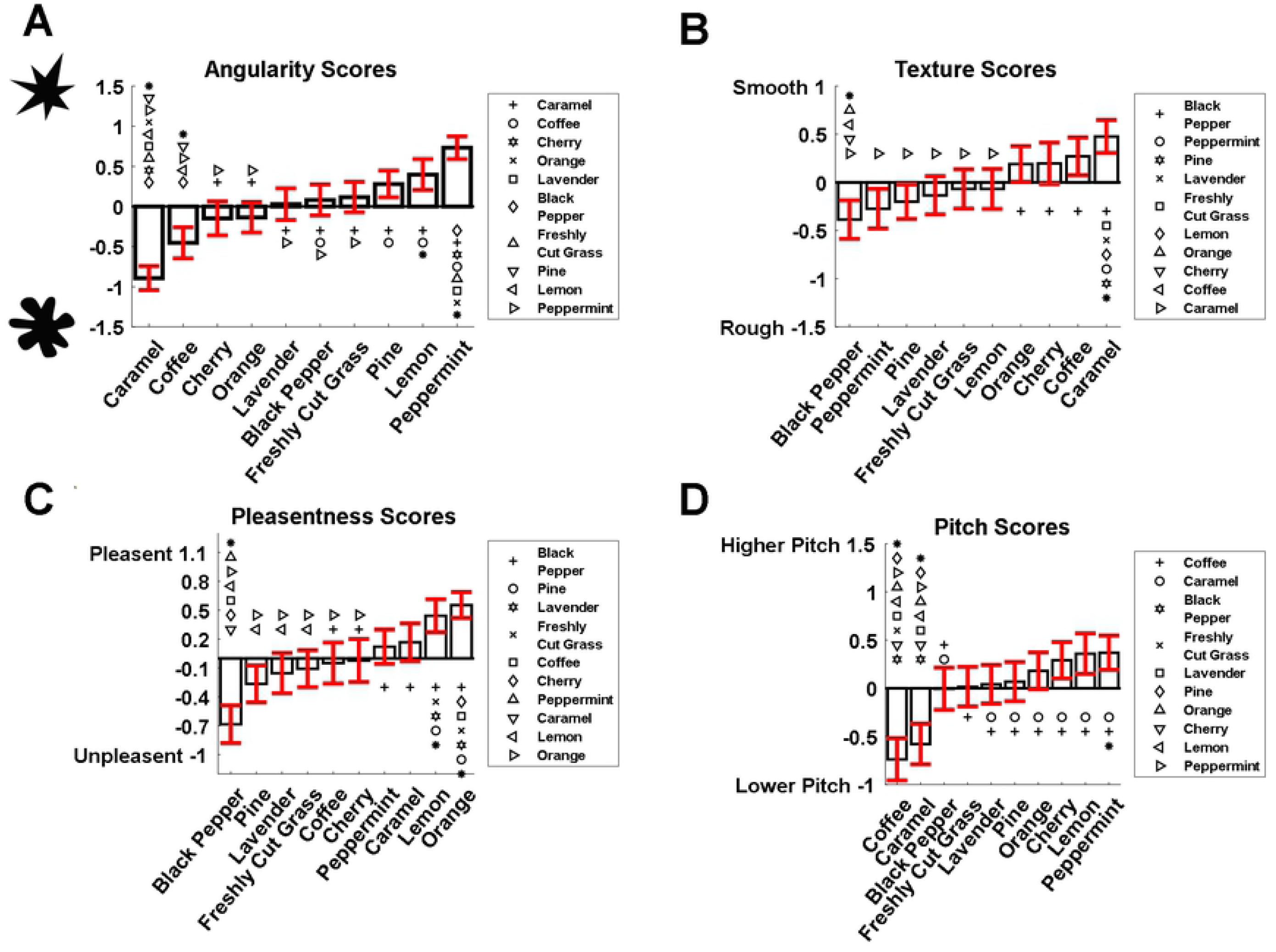
Shape, texture, pleasantness and pitch scores. (A – C) Mean scores for the 10 odours after z-score normalisation using the grand mean, asterisks mark the odours that are significantly different from the scale’s original grand mean. Errors bars show a 95% confidence interval of the respective odour. The legend shows markers indicating which odours have significantly different means from the respective odour (e.g., the significantly different odours from coffee in (**A**) are lemon, peppermint and pine. (D) Shows the same information as (A – C) apart from the mean value used to calculate it z-score is that of log2 of the original ratings.

### Genre and emotions

To assess if odours affected the genre and emotion selections chi-squared tests of independence were conducted. This revealed that the odours impact both the choice of genre (*x*^2^ = 138.20, *P* < 0.05) and participants emotional response (*x*^2^ = 187.54, *P* < 0.05). Consequently, chi-squared tests for goodness of fit were conducted to see which of the presented stimuli were significantly different from a chance selection. The odours that were significantly different from a chance selection in the genre association task are black pepper, caramel, cherry, coffee, freshly cut grass, lemon and orange (*P* < 0.05) (See Fig 2A). The odours were significantly different from chance selection in the emotion association task (*P* < 0.05) (See Fig 2B).

**Fig 2.**
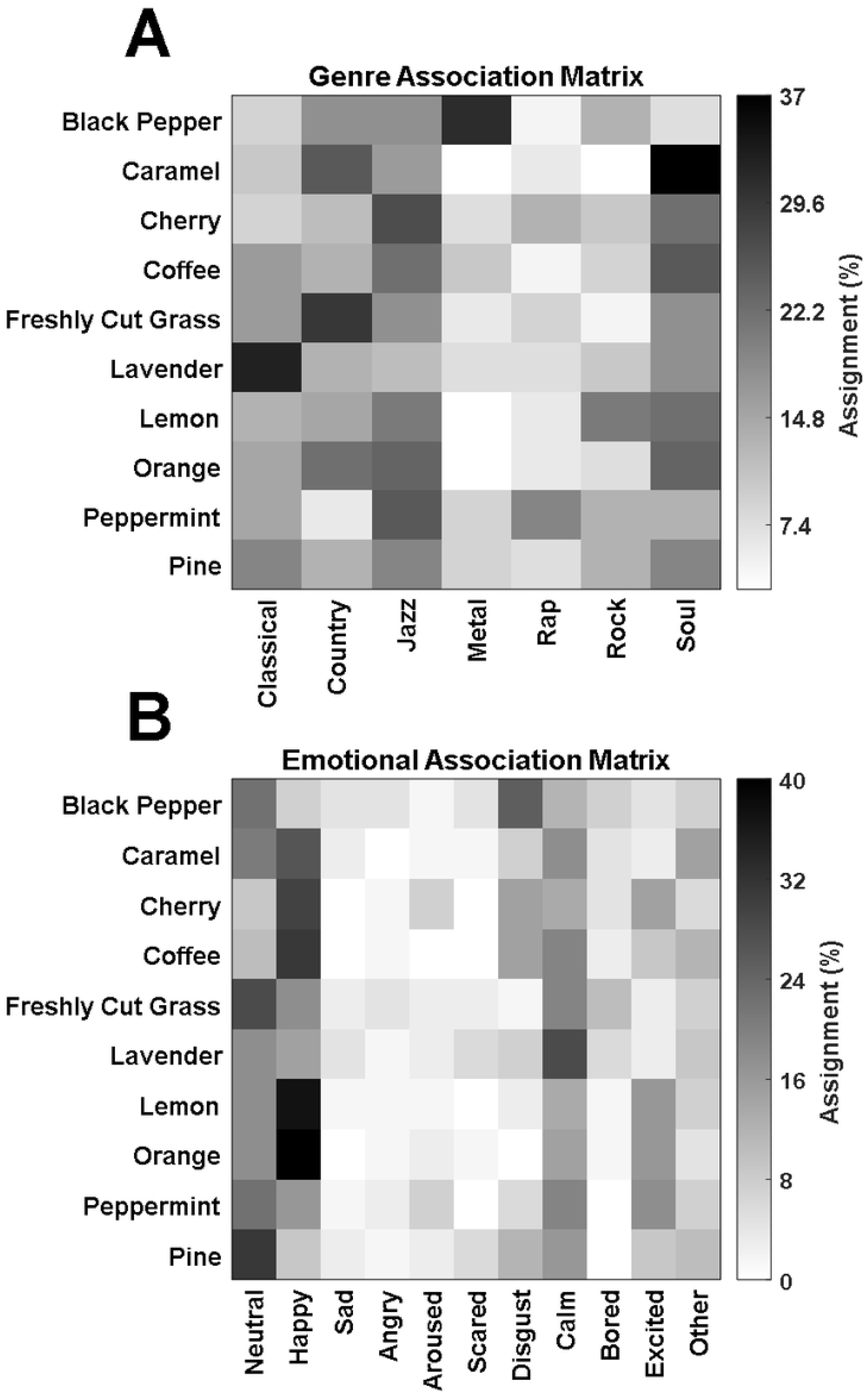
Musical genre and emotional association matrices. (A) Association matrix between the 10 odours and the 11 possible emotional selections. (B) Association matrix between the 10 odours and the 7 genres.

### Colours

Due to many possible colour choices, 343 interpolated colours from L*a*b* colour space were chosen (See Fig 3A). The colour the user had selected was then mapped to one of the 343 colours based on the lowest delta E 2000 error to determine the perceptually closest colour. This was consequently used as the representative selection, shown in Fig 3B. The median hue angle for the commonly selected colours is shown in, Fig 3C. Each participant only reported one colour for each odour. A one-way t-tests (test value = 50) was conducted with a Bonferroni corrected alpha of 0.005 (0.05 / 10) to determine if the selected lightness values were significantly different from the range’s midpoint and default slice of 50. The significantly different odours are; (where *L\** represents the mean lightness score from the L*a*b* colour space) caramel (*P* < 0.005, *t* = 4.30, *L* =* 59.78), cherry (*P* < 0.005, *t* = 3.66, *L* =* 58.14), coffee (*P* < 0.005, *t* = -2.18, *L* =* 44.73), freshly cut grass (*P* < 0.005, *t* = 3.47, *L* =* 56.91), lavender (*P* < 0.005, *t* = 4.82, *L* =* 60.11), lemon (*P* < 0.005, *t* = 15.10, *L* =* 76.51), orange (*P* < 0.005, *t* = 12.53, *L* =* 70.10), peppermint (*P* < 0.005, *t* = 7.5, *L* =* 66.35) and pine (*P* < 0.005, *t* = 3.71, *L* =* 58.68). A chi-square test of independence was conducted on the hue angles of the colours. Due to the large number of possible angles binning was implemented (*N* = 15), a chi-square test for goodness of fit was conducted on the binned hue angles to see if the colours selected differ from chance selection, this revealed the colour selections for each odour significantly differ from chance selection (*P* < 0.005).

**Fig 3.**
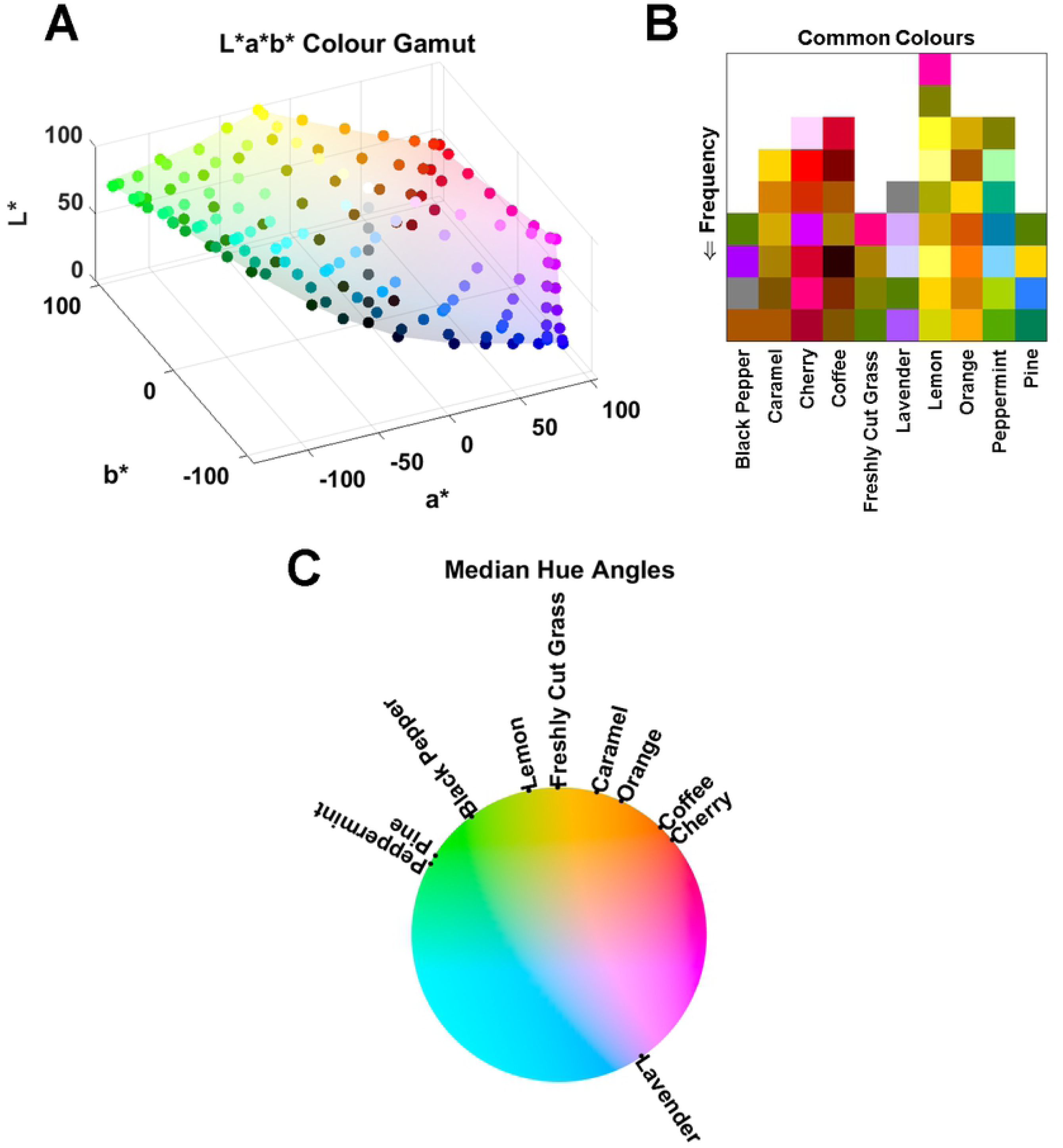
Commonly selected colours and median hue angles. (A) L*a*b* colour gamut showing the interpolated points used to determine the perceptually closest colour. (B) Common colours selected by the participants where each colour has been mapped more than twice. (C) Cylindrical representation of the L*a*b* colour space showing the median hue angle of the commonly selected hues for each odour

### Identification dependencies

To determine if the proportions (correct and incorrect) where significantly different, chi-squared tests for goodness of fit where conducted this revealed that the proportions for black pepper (*x*^2^ = 85.76, *P* < 0.05), caramel (*x*^2^ = 47.05, *P* < 0.05), coffee (*x*^2^ = 7.52, *P* < 0.05), freshly cut grass (*x*^2^ = 19.88, *P* < 0.05), lemon (*x*^2^ = 51.88, *P* < 0.05), orange (*x*^2^ = 7.52, *P* < 0.05), peppermint (*x*^2^ = 56.94, *P* < 0.05), and pine (*x*^2^ =,73.52 *P* < 0.05) varied significantly. To access if the variation between correct and incorrect identification was statically significantly a two-way ANOVA was conducted for each odour. This revealed that there was no significant variation for the angularity ratings (see Fig 4A), the significant odours for the smoothness ratings are lavender (*P* < 0.05) and caramel (*P* < 0.05) (see Fig 4B). The significant odours for the pleasantness ratings are lavender (*P* < 0.05) and peppermint (*P* < 0.05) (see Fig 4C) with the latter also being significant in the pitch ratings (see Fig 4D).

**Fig 4.**
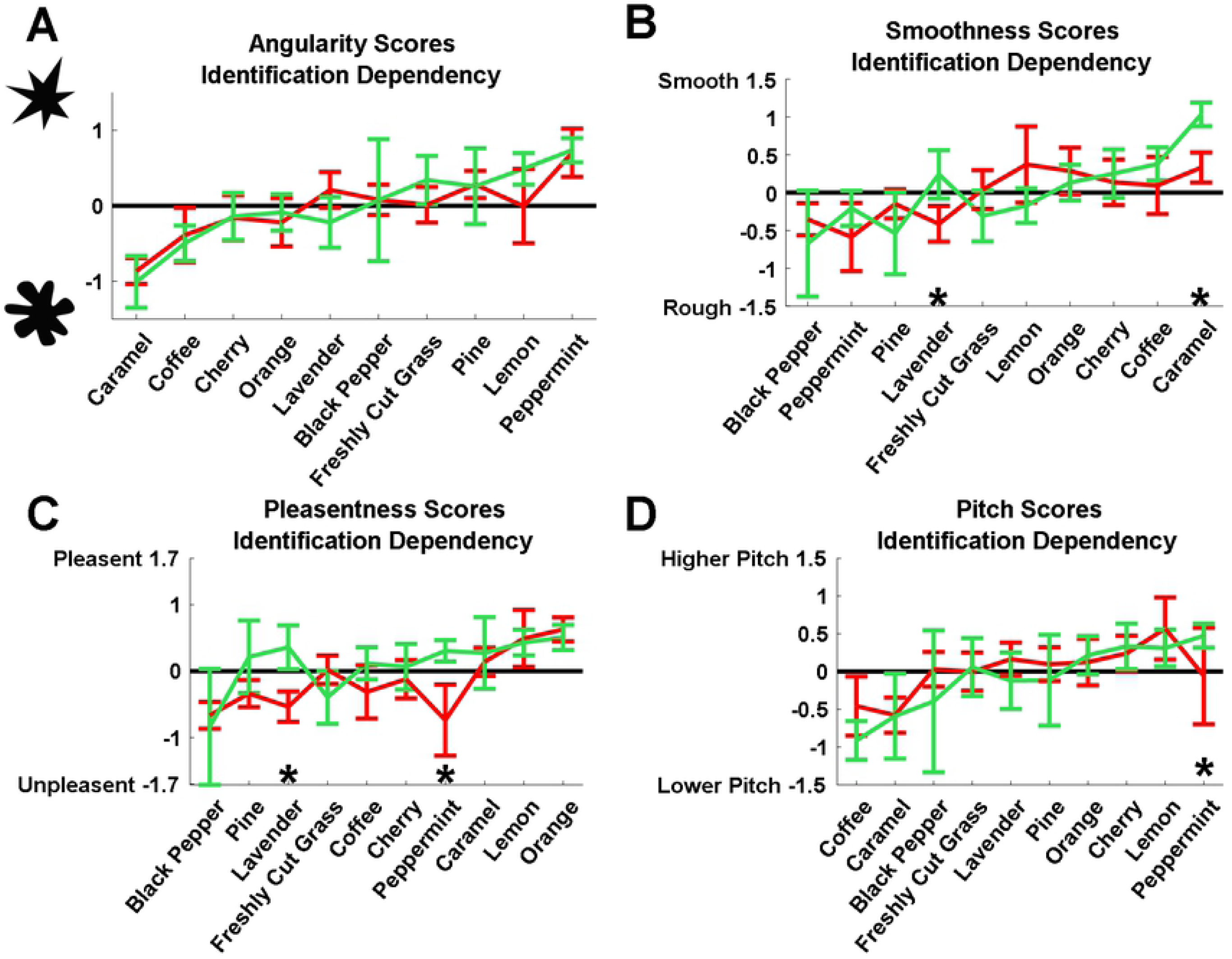
Angularity, smoothness, pleasantness and pitch identification dependencies. Mean z-scores asterisks detonate odours where there is significant variation between the correct and incorrect ratings. The green markers detonate correct classification and the red markers detonate incorrect classification. The error bars show a 95% confidence interval.

To access if explicit knowledge of the odour affected the emotional and genre dimensions the relative difference between correct and incorrect identification was calculated for each odour. Fig 5A shows that the knowledge of an odour does affect the emotional dimensions and has Frobenius norm of 323.06. Peppermint, for example, was perceived as less happy and angrier. Generally, the odours were perceived as being more neutral, calming and slightly disgusting. From Fig 5B we can see that knowledge of the odours affects the genre dimensions, for example, peppermint is perceived to be less jazz and more metal and rap the genre association matrix has a Frobenius norm of 339.70.

**Fig 5.**
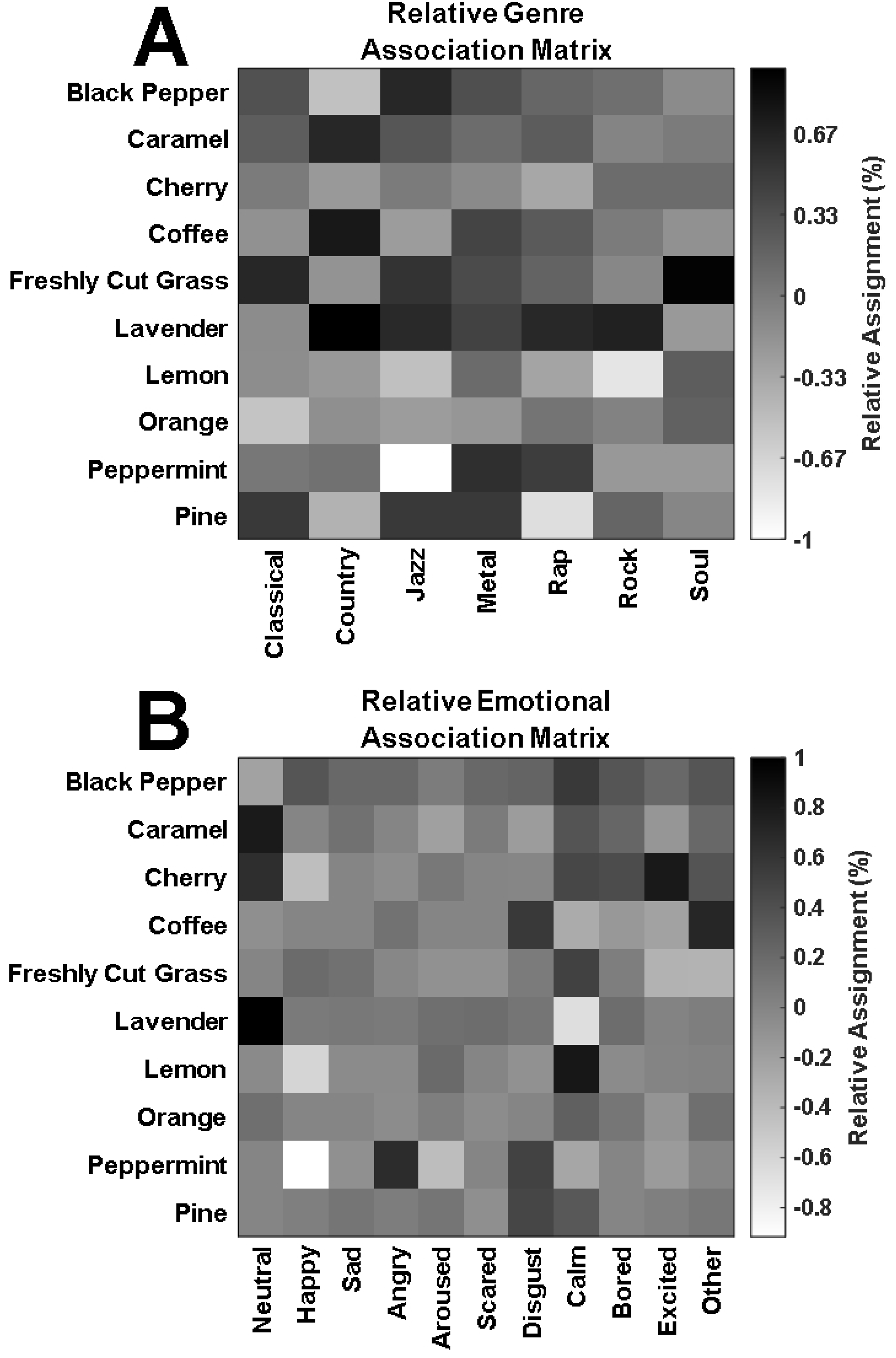
Relative musical genre and emotional association matrices. (A) Relative association matrix between the 10 odours and the 7 genres. (B) Association matrix between the 10 odours and the 11 possible emotional selections.

Following the same procedure as the colour analysis above, common colours for the misclassified odours are shown in Fig 6. To determine if the observed proportions between the common colours and the common misclassified colours were significantly different, chi-squared tests for goodness of fit where conducted this revealed that the proportions for cherry (*x*^2^ = 14.00, *P* < 0.05), coffee (*x*^2^ = 7.77, *P* < 0.05), lavender (*x*^2^ = 4.28, *P* < 0.05), lemon (*x*^2^ = 18.00, *P* < 0.05), orange (*x*^2^ = 10.50, *P* < 0.05) and peppermint (*x*^2^ = 14.00, *P* < 0.05) are significantly different. The same pattern is observed for correct and incorrectly identified odours, which is consistent with the idea that crossmodal correspondences between olfaction and colour do not rely on explicit knowledge of the odour’s identity.

**Fig 6.**
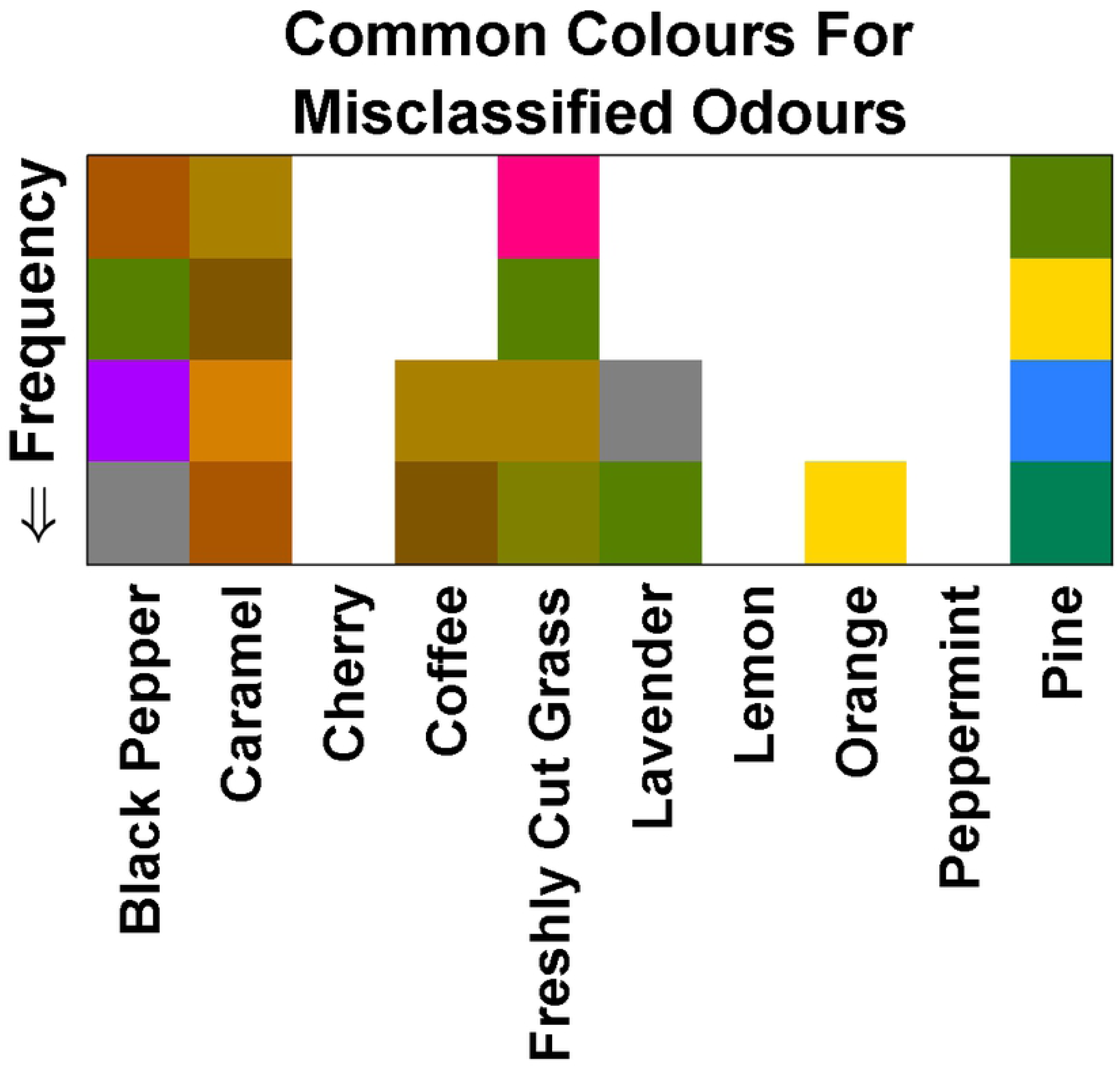
Common colours for misclassified odours. Common colour selections where each colour has been incorrectly classified and been mapped more than twice.

### Identification

The task of participant was to identify the given odour by selecting one of the 25 possible odours in the list. Retrospectively, a twofold identification was considered, exact identification and categorical identification. Exact identification was achieved 45.74% of the time by correctly identifying the current odour, participants could select an odour label from a compiled list of 25 different odours. The top three correctly identified odours are peppermint (84.21%), lemon (80.88%) and orange (63.16%). The top three misclassified odours are black pepper (10.29%), pine (12.24%) and caramel (20.59%) (see Fig 7). Retrospective category identification was determined by the participants’ ability to pick another odour in the same category following the fragrance classes outlined in [35]. An accuracy rating of 62.94% was achieved for category identification, each potential classification belonged to only one category. A Pearson correlation indicated that there was no strong correlation between the age of the participant and their identification accuracy (*P* = -0.17).

**Fig 7.**
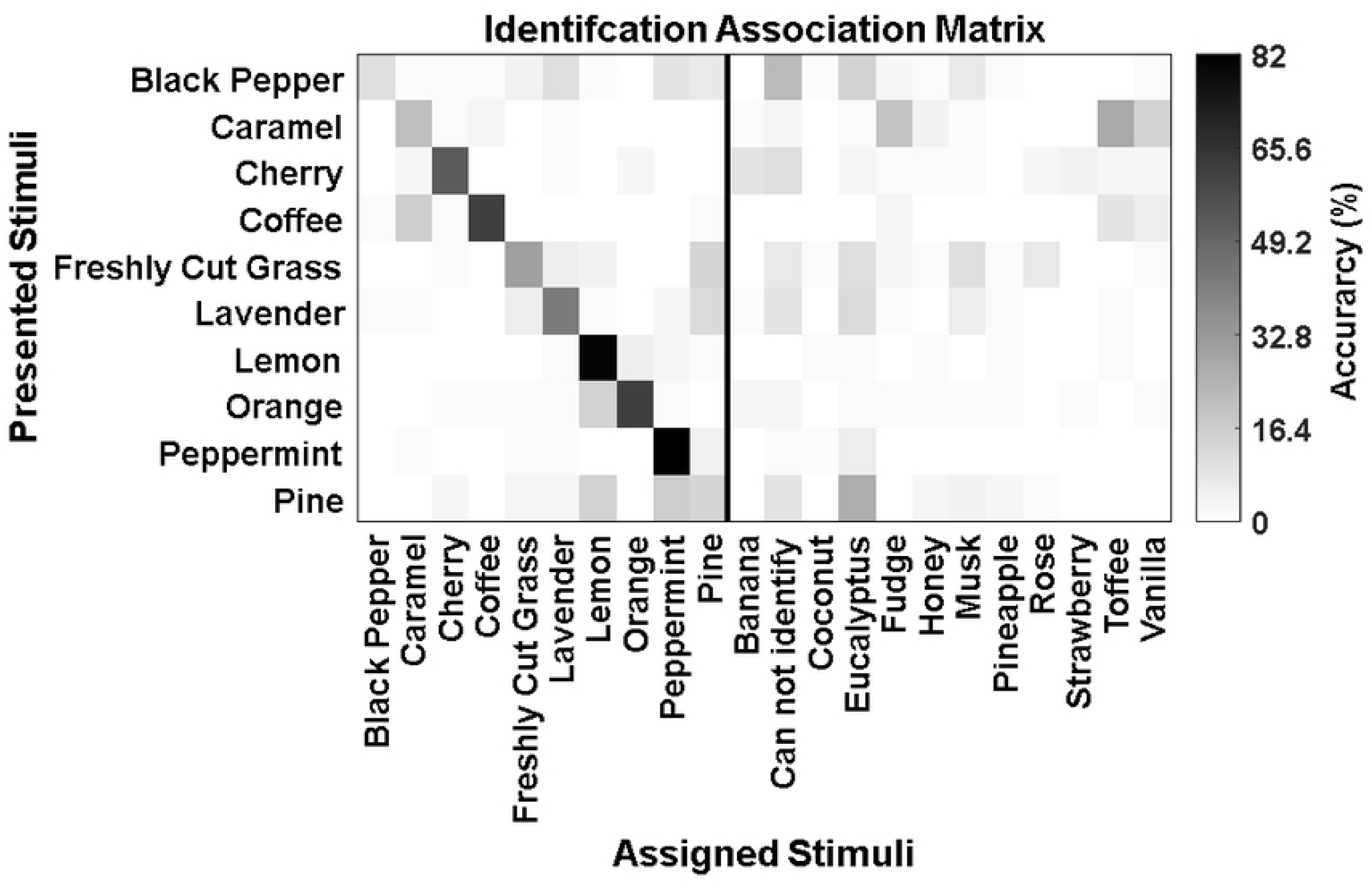
Identification association matrix. Association matrix for the 10 odours, along with an additional 12 misclassifications.

### Principal component analysis

Factor analysis using Principal Component Analysis (PCA) was conducted on; the shape ratings, texture ratings, pleasantness ratings, pitch ratings, identification accuracy, the colour dimension (lightness), the emotional dimensions and the genre dimensions. Due to the ratings and dimensions being on different scales z-score normalisation using the odour-wise mean and the standard deviation on the original dataset before PCA. Based on inspection of the score plot, four principal components were kept, explaining 81.57% of the total variance and have an eigenvalue of at least 1. The principal components 1 through 4 explain 32.49%, 25.60%, 14.43% and 9.04% of the total variance respectively. The first two principal components are shown in Fig 8A with the loading matrix, shown in Fig 8B.

**Fig 8.**
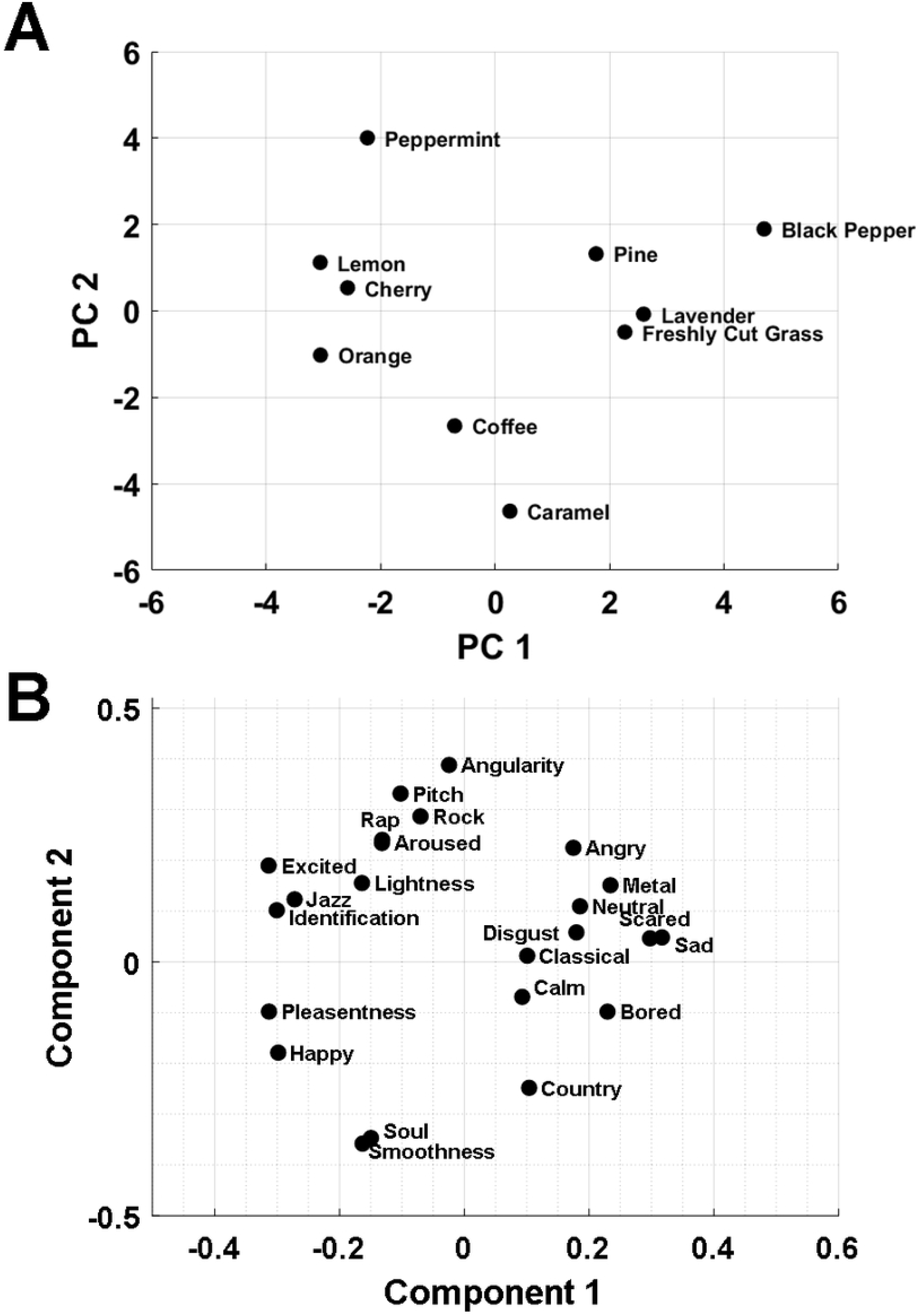
PCA score and loadings. (A) Score plot of the perceptual ratings for each odour. (B) Loading’s plot showing the covariance coefficients for each of the dimensions used in the PCA.

## Discussion

Our results further our understanding of crossmodal processing and how olfactory sensations interact with the perception of angularity of shapes, smoothness of textures, perceived pleasantness, pitch, genre and emotions. Our first hypothesis was that robust associations exist for common aromatic compounds and we find evidence for such associations, consistent with prior findings; olfactory associations between the angularity of shapes [20], smoothness of texture [36] and pitch [12] suggesting that the hedonic qualities are important factors in crossmodal correspondences. Our second hypothesis was that knowledge of a odours identity would affect some of the reported associations, consistent with the findings reported by [18] between odours and colours and angularity of shapes. Odours produce reliable and distinct colour profiles, which are more consistent with the explicit knowledge of an odour’s identity. Furthering these findings, we show the reliability and extent of these associations, and that the knowledge of an odour’s identity partly affects their reported associations between the other modalities with the exemption of the angularity of shapes.

The PCA analysis score plot shows the perceptual similarity between the olfactory stimuli, for example, (lemon, cherry and orange), (lavender, freshly cut grass and pine), (coffee and caramel) obtained similar results in most, but potentially not all, ratings analysed using PCA. The PCA loadings plot suggests that the hedonic values are a strongly influenceable factor, that is, the strong loadings of the pleasant (i.e., happy, excited and calm), unpleasant (i.e., sad, angry and scared) and the ‘pleasantness dimension’ are at least moderately associated to the other dimensions reported in this paper. Additionally, the loadings plot shows strong associations between the angularity of shapes, textures, pitch, emotional and genre dimensions. Odours judged to be the rounded shape tend to be associated with a smooth texture, be lower in pitch, be more soul. Whereas, the angular shape appears to be strongly associated with the genre dimension rock. Moderate associations exist between the angularity of shapes with the angular shape being associated with lighter colours, being more rap, angry, arousing and exciting. Strong associations exist between the odours and the smoothness ratings with smooth being perceived as more soul, happier and less angry. Moderate relationships exist with smooth being perceived as more pleasant, lower in pitch and being more country. With the rough texture being associated to be more metal, rock, neutral, sad and angry. The strong associations between pleasant odours are being less metal, disgusted and being happier. With moderate relations being more excited, soul, less sad, easier to identify with an increase in lightness. Higher pitches are strongly associated with being lighter in colour and being more rock with moderate relations to being more exciting, arousing, rap, and being less soul. Darker colours are moderately associated with being more metal and disgusted while lighter colours are moderately associated with being more rock and exciting. The strong associations between the odours and identification rate are being more happy, exciting and being less sad and disgusting. Moderate relationships exist between a higher identification rate and being more jazz and less metal, classical, neutral, disgusting and boring. All relationships uncovered in the PCA analysis can also be applied in reverse (i.e., the odours perceived as being smoother are generally assigned to being less angular and more rounded in shape). Consistent with the findings from [37], The PCA shows strong relationships towards the emotional and musical dimensions, for example, the relationships between the emotion ‘happy’ are being less more jazz and soul and less metal. The relationships between ‘angry’ are being more metal and less soul.

Future work stemming from these findings could include assessing if congruent vs incongruent associations defer any advantages in an objective task. The moderate relations discovered in the PCA analysis, between odours and the different relationships could be explored further with a more tailored set of experiments. The cross-model mediations (i.e., the intercorrelated dimensions) uncovered during the PCA may be explored further to uncover the robustness of the associations. Additional sensory dimensions could be added, gustatory could be explored, towards the goal of an olfactory psychophysical framework, to aid in a variety of different applications ranging from olfaction enhanced multimedia to marketing. Due to the nature of olfaction, it may be the case that taste played a minor role in the associations reported in this paper. For example, [38] reported the bitter tastes were associated with angular shapes, whereas, sweet tastes are associated with a more rounded shape. In contrast with a later study, [20] reported a strong association between the angularity of shapes and the perceived sourness/bitterness of odours.

## Author contributions

RJW and SMW conceived and designed the experiments. RJW performed the experiments. RJW and SMW analysed the data. AM and SMW contributed reagents. RJW programmed the analysis tools. SMW and AM provided supervision. RJW, SMW and AM wrote the paper.

